# RecurSieve: A Python Package to Detect Subtle Gene Expression Differences Across Single-Cell RNA-seq Comparisons Using Iterative Machine Learning

**DOI:** 10.64898/2026.11.24.768193

**Authors:** Anushi Mittal, Gregory Chin, Connor Frankston, Lamya Ben Ameur, Arpiar Saunders, Kenny Pavan

## Abstract

Most existing methods for performing differential expression testing using single-cell RNA-seq data evaluate genes independently and primarily detect shifts in mean RNA levels. These methods may thus miss RNAs that distinguish experimental groups jointly or through changes in the shape of genes’ expression distributions. To address this limitation, we developed RecurSieve, an iterative ensemble learning method that combines shared expression classification with unsupervised clustering to recursively identify gene sets with group-based expression differences. Because experimental labels apply to heterogeneous populations of single cells within each comparative group, RecurSieve allows dynamic label reassignment during iterative training to detect subtle, non-linear expression patterns. In benchmarking experiments using simulated data with staged distributional expression differences, RecurSieve recovered more genes (2.3-3.9 of 10 staged genes) across groups as compared to standard methods (Wilcoxon, t-test, and logistic regression; 0.0-0.7 staged genes). When applied to a single nuclei RNA-seq analysis containing low and high-pathology stages of Alzheimer’s disease (46 human donor brains), RecurSieve extended results from standard approaches by identifying genes including *WWOX*, *DPP10*, *KCNIP4*, *HS6ST3*, and *CNTN5* which tend to exhibit stage-specific distributional expression differences accompanied by only minor changes in average expression. A subset of the genes exclusively detected in RecurSieve analysis had been previously associated with Alzheimer’s disease, underscoring the potential importance resulting from the unique sensitivity of our method. These simulated and empirical single-cell RNA-seq results demonstrate how the RecurSieve framework complements standard methods for identifying genes with unique group-specific expression differences independent of changes in average expression.

**Availability and Implementation:** RecurSieve is an open-source and pip-installable package that can be found at the repository https://github.com/ArpiarSaundersLab/RecurSieve along with documentation and usage tutorials.

## 1. Introduction

Single-cell transcriptomics has become a central tool for studying the molecular properties of complex tissue, including identifying candidate therapeutic targets in disease contexts^1–4^. However, extracting biological insight from these large and high-dimensional datasets remains challenging. Differential expression (DE) analysis is among the most foundational and frequently used approaches to characterize single-cell RNA-seq (scRNA-seq) datasets, owing to its capacity to identify statistically significant changes in gene-specific mRNA levels^5^. Tools such as DESeq2^6^, edgeR^7^, and Scanpy^8^ are widely used in bulk and scRNA-seq DE workflows, and common statistical tests include the Wilcoxon rank-sum test^9,10^, the t-test^11,12^, and logistic regression^13,14^. The Wilcoxon test, for example, is a standard choice due to its robustness to outliers and lack of reliance on assumptions of normality.

Despite their established utility, standard DE methods may fail to detect important differences in RNA changes across groups. For example, they may miss genes that jointly distinguish disease states or that differ in the shape of their expression distributions, changes relevant to many biological processes^15,16^. These limitations arise because established DE methods evaluate each gene independently, leaving co-regulation to other methods or strategies explicitly focused on identifying regulatory networks, or use univariate tests that primarily detect shifts in central tendency (mean or median). Consequently, these standard DE tests are largely insensitive to patterns where distribution shape changes - such as bimodality or skewness - yet leave mean RNA levels similar. Such subtle shifts may represent critical transcriptional transitions in cell state populations that are largely invisible to standard ranking methods^16,17^. Current strategies to address this include pseudobulk aggregation^18^, gene set (module scoring)^19^, and multivariate machine learning approaches^20^. However, balancing interpretability with robustness to technical noise remains challenging. There remains a need for methods that explicitly model joint expression relationships and distributional shapes without reliance on pre-defined gene sets.

Additionally, standard scRNA-seq DE methods compare experimental groups by assigning static labels to all cells of each group, in spite of the fact that groups often consist of molecularly heterogeneous cell populations. DE analysis based on static labels could mask detection of more subtle biological processes - such as disease states - that vary more in frequency than quality across experimental groups^21,22^. Indeed, quantitative pathology of brain tissue from 84 aged donors spanning the Alzheimer’s disease continuum showed that donors with low pathology can harbor cells in disease-relevant transcriptional states^23^. In contrast to per-gene DE testing, feature selection identifies sets of genes that jointly discriminate groups, making it naturally suited to detecting coordinated and distributional changes. Feature selection methods that allow dynamic, data-driven swapping of group labels across cells could help address this effect and sensitivity for detecting changes in RNA levels forecast by initial group assignments^17^. Recently, state-based approaches have relaxed static label assignments by estimating group association at the level of individual cells or cell neighborhoods^24,25^. However, these methods primarily test for differential abundance of cell states rather than identifying the genes that distinguish them.

In this study, we developed RecurSieve, a method for scRNA-seq feature selection that is not bound by assumptions of univariate RNA level changes, gene independence, or static cell group assignments and thus sensitive to discover a more complete set of genes with complex RNA level changes across experimental groups. RecurSieve implements a hybrid recursive framework combining a Random Forest (RF) classifier^26^, Gaussian Mixture Model^27^ (GMM) based clustering, and dynamic cell label group assignment. We tested RecurSieve’s expanded sensitivity by first analyzing synthetic benchmark data featuring staged genes with controlled distributional shifts (bimodal, trimodal, and skew), followed by empirical analysis of a scRNA-seq Alzheimer’s disease (AD) dataset spanning multiple disease stages^28^. In synthetic experiments, RecurSieve recovered more staged gene changes than standard DE methods. Analysis of Alzheimer’s disease data revealed stage-associated changes in gene-specific RNA levels distributions with inconspicuous mean expression differences. These results suggest RecurSieve analysis offers an interpretable strategy for discovering biologically meaningful alterations in RNA levels that are often overlooked by standard univariate DE testing methods.

## 2. Results

### 2.1 RecurSieve Iterative Workflow Converges on Stable Gene Panels

The RecurSieve algorithm uses a closed-loop system to iteratively update population cell labels associated with each compared population using a semi-supervised ensemble approach, collapsing genes into a cumulative expression axis at each step (**Figure 1A**). This workflow initiates with standard preprocessing (quality control, normalization, log-transformation, and highly variable gene selection) followed by an initial RF gene feature classification based on the initial user-defined experimental labels.

**Figure 1:**
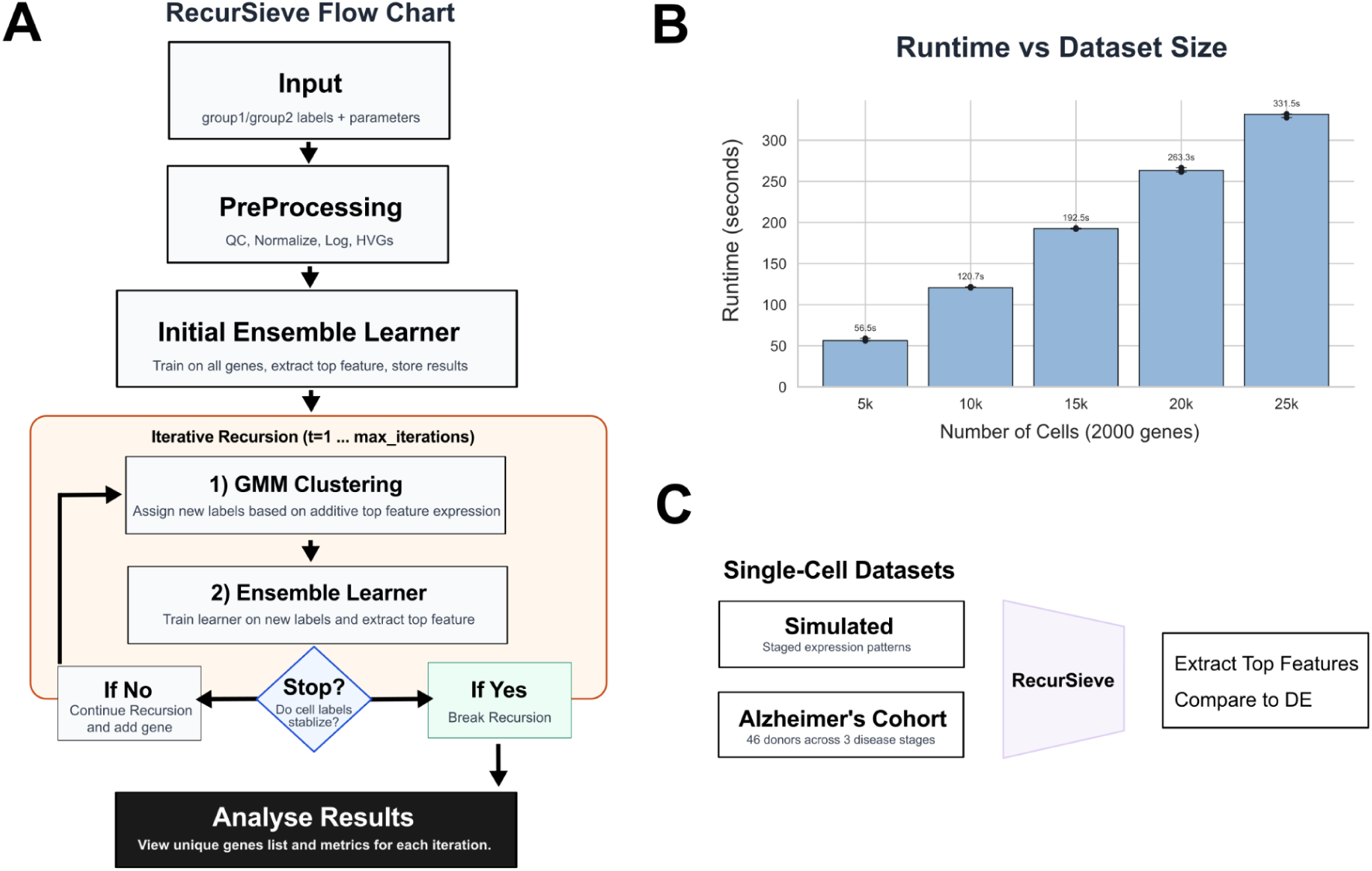
RecurSieve Overview, Runtime, and Assessment. **(A)** Schematic of the RecurSieve algorithm. The pipeline begins with preprocessing (quality control, normalization, log-transformation, and selection of highly variable genes), followed by iterative recursion where an ensemble classifier identifies top features, Gaussian Mixture Modeling (GMM) generates dynamic labels, and recursion continues until label stabilization. (B) Median runtime performance of RecurSieve as a function of dataset size. Execution time was measured for datasets containing 2,000 highly variable genes with increasing cell counts (5k-25k cells). Error bars show min-max range of *n=3* trials. This code was run on a MacBook with an 8-core CPU, 16 GB RAM, running macOS Sequoia 15.7.8. **(C)** Experimental validation strategy. RecurSieve performance was assessed using two distinct datasets: a simulated dataset generated with Splatter to introduce known staged, non-linear gene expression patterns across groups, and a real-world single-nuclei RNA sequencing dataset from an Alzheimer’s disease cohort (n=46).

From this initial model, RecurSieve extracts the top-ranking feature (gene) based on Gini importance^29^. This gene, combined with the total unique molecular identifier (UMI) counts of each cell (normalized + log1p-scaled), is used to fit a two-component Gaussian Mixture Model (GMM), which generates dynamic training labels for the next iteration. The algorithm then enters a recursive cycle: the new labels redefine the ground truth, allowing the RF to identify the next most informative gene. This newly selected gene is added to the cumulative panel, and the GMM is refitted using the aggregated expression of the entire panel plus library sizes (Default: mean). This process repeats until label assignments stabilize, defined as a label disagreement rate below 0.01 between consecutive iterations, or no improvement is made in label stabilization (Default: 10 iterations), with runtimes being tractable to modern single-cell dataset sizes (**Figure 1B**).

To validate the efficacy of this iterative framework, we applied RecurSieve to two distinct datasets: a simulated dataset engineered with staged, non-linear gene expression patterns (bimodal, trimodal, skew, and coexpression) to benchmark recovery rates against standard differential expression methods, and a single-nuclei RNA sequencing dataset from an Alzheimer’s disease cohort^28^ comprising 46 donors across three disease stages (**Figure 1C**).

### 2.2 RecurSieve Recovers Non-Linear and Staged Gene Signatures in Simulated Data

To test RecurSieve’s sensitivity to non-linear expression patterns, we generated synthetic single-cell datasets using Splatter^30^ (2,000 cells, 2,000 genes) and embedded ten staged genes with complex distributional shifts. These scenarios included bimodal, trimodal, skewed, and co-expression patterns where the underlying mean and median expression remained constant between groups, mimicking biological situations where disease states alter gene regulation variability or cell subpopulation ratios without changing the overall average abundance^16,31^. We benchmarked RecurSieve against standard DE methods implemented in Scanpy^8^ (Wilcoxon, t-test, logistic regression) to determine which approach could successfully recover these subtle, shape-based signals.

We evaluated five distinct scenarios: true null (shuffled labels), bimodal, trimodal, skewed, and co-expression. In the bimodal, trimodal, and skewed scenarios, the staged genes in Group 2 were engineered to have multiple peaks or asymmetric tails that closely balance the central tendency of the staged genes in Group 1. As anticipated, standard DE methods largely failed to detect the staged genes across these scenarios. The Wilcoxon, t-test, and logistic regression approaches recovered an average of 0.13 (mean recovered/ total conditions) genes per trial (n=25) with seed stability testing (seeds=25) across these conditions (**Figure 2A**). Non-zero recovery included the t-test bimodal test, which achieved a mean of 0.3 genes recovered across all trials (SEM±0.21). Additionally, logistic regression recovered 0.1, 0.1, 0.3, and 0.7 genes for bimodal, trimodal, skewed, and co-expression scenarios, respectively (SEM: ±0.06; ±0.06; ±0.11; ±0.17). These results confirm that when the central tendency is conserved, univariate tests relying on location shifts are largely blind to the distributional changes we tested.

**Figure 2:**
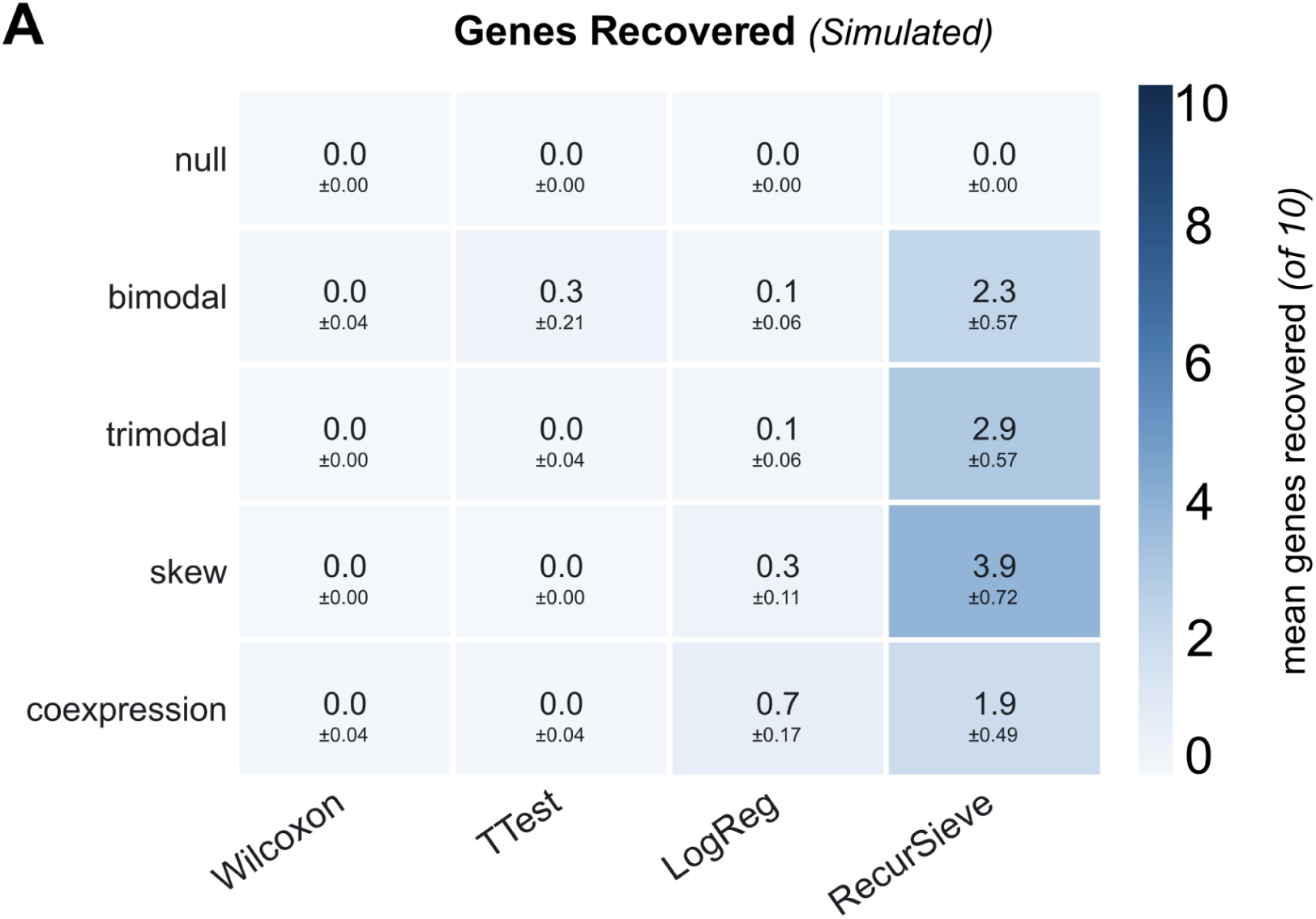
Synthetic Dataset Testing. **(A)** Simulation benchmark showing mean number of staged genes recovered (out of 10) over 25 simulated dataset trials, with 25 seed stability repetitions. RecurSieve outperforms standard differential expression (DE) methods (Wilcoxon, t-test, logistic regression) in recovering bimodal, trimodal, skew, and coexpression patterns in simulations. Values are mean ± SEM across the 25 dataset means. Wilcoxon and t-test genes were filtered at Benjamini–Hochberg adjusted p < 0.05. Logistic regression does not produce p-values, so its top 50 genes by score were used. RecurSieve does not filter on p-values.

When examining the synthetic results of the 25 trials, RecurSieve recovered 2.3 genes (mean recovered of 10) in the bimodal scenario (SEM±0.57), 2.9 in the trimodal scenario (SEM±0.57), and 3.9 genes in the skew scenario (SEM±0.72) (**Figure 2A**). Notably, in the co-expression benchmark where genes exhibited identical marginal distributions but became positively correlated only within Group 2, RecurSieve recovered a signal of 1.9 genes (SEM±0.49), while all other tested DE methods recovered fewer genes (≤0.7). Additionally, in the true null scenario where labels were shuffled and no biological signal existed, RecurSieve correctly identified 0.0 genes, indicating that iterative label transfer did not generate spurious signal when no structured difference was present across the initial experimental groups. These results suggest RecurSieve is sensitive to detect complex, non-linear gene expression relationships and dependency structures across comparative groups for which standard DE testing is insensitive.

### 2.3 Gene Signatures Found in Single-Cell Alzheimer’s Dataset

To evaluate RecurSieve on empirical data, we analyzed a single-nuclei Alzheimer’s disease dataset^28^ comprising 424,528 nuclei from 46 donors across three cortical regions (prefrontal cortex, precuneus, and primary visual cortex), with comprehensive metadata (**Methods 4.2**). Using the ‘Disease.Group’ annotation and all cell types combined (Excitatory, Inhibitory, Astrocytes, Microglia, Oligodendrocytes, OPCs, and Vascular), we applied RecurSieve to distinguish features between low and high disease pathology groups in all cell types and Excitatory neurons (**Figure 3, 4**), while maintaining default preprocessing parameters (**Methods 4.2.1**). For all cell types, the algorithm converged after 41 iterations, achieving label stability in the GMM step (<1% delta) (**Methods 4.1.3**). For comparison, we performed standard Wilcoxon-based differential expression analysis using Scanpy and conventional thresholds^32^ (p < 0.05; |log2FC| ≥ 0.1).

**Figure 3:**
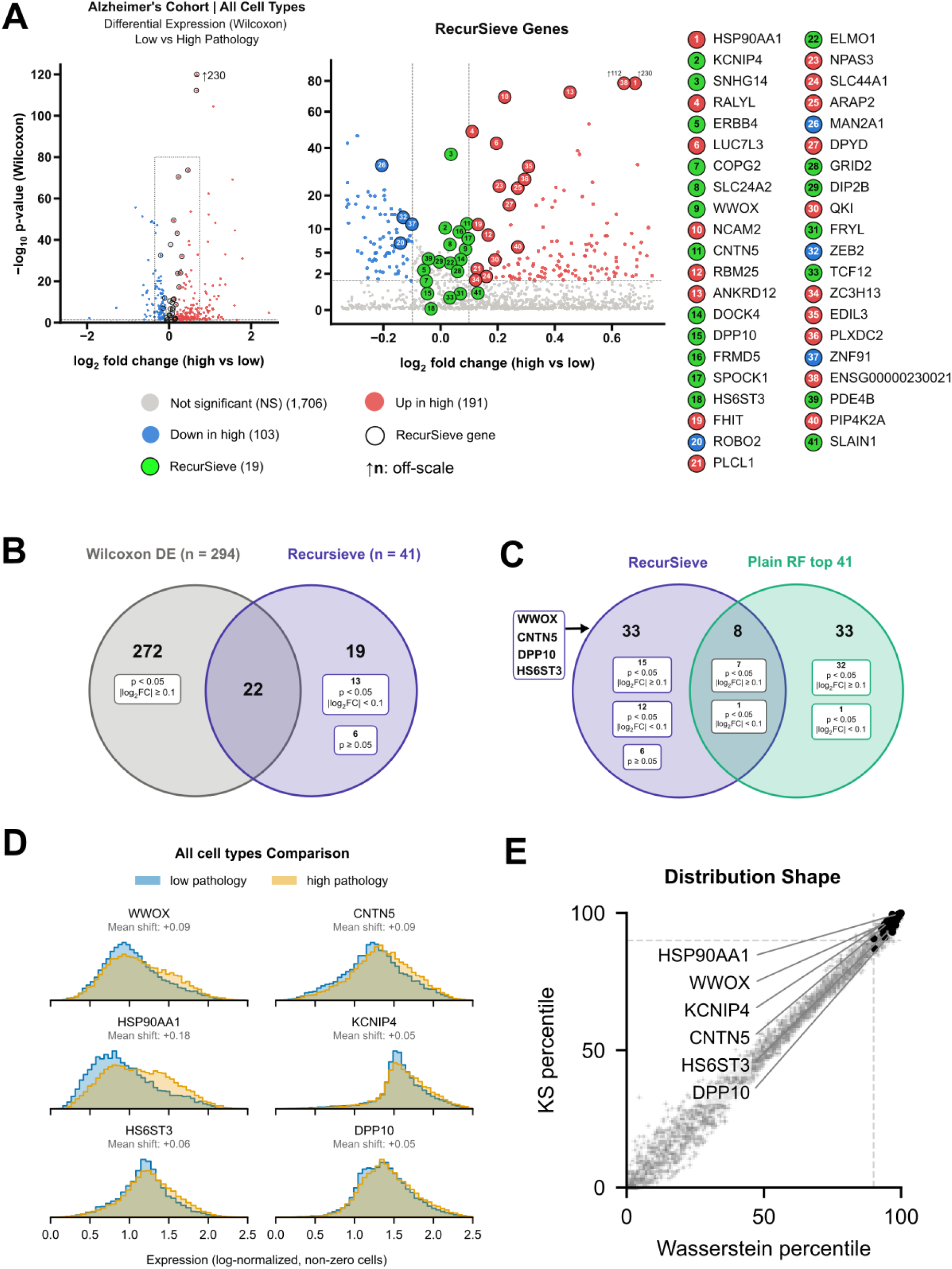
All Cell Types Combined Results in Alzheimer’s Single-cell Data. **(A)** Volcano plot comparing RecurSieve-selected genes (n=41) versus DE-identified genes (Wilcoxon test) in an Alzheimer’s disease single-nuclei cohort (All Cell Types). RecurSieve reveals genes with lower fold-changes, capturing expression shifts missed by DE. (Right) ranked list of the 41 RecurSieve-identified genes in the Alzheimer’s cohort, ordered by iteration of selection. **(B)** Venn diagram illustrating the overlap between genes identified by standard Wilcoxon differential expression (DE) analysis (n=294; p < 0.05, |log₂FC| ≥ 0.1) and the RecurSieve algorithm (n=41) in all-cell-types. Of the 41 genes selected by RecurSieve, 22 were also detected by DE, while 19 were found to be RecurSieve genes where 13 had p < 0.05 but fell below the log₂FC threshold (|log₂FC| < 0.1), and 6 did not reach statistical significance in the DE analysis (p ≥ 0.05), highlighting RecurSieve’s ability to capture subtle, coordinated expression patterns missed by univariate thresholds. **(C)** Top 41 RecurSieve genes for the all-cell-types comparison displayed in a Venn diagram illustrating the overlap between genes identified by RecurSieve vs. standard RF feature importance. (33) belonging to RecurSieve, (33) belonging to RF, and (8) intersecting both. RecurSieve recovered genes include *WWOX*, *CNTN5*, *DPP10*, and *HS6ST3*. **(D)** Expression distributions (non-zero) and mean shifts of *WWOX*, *CNTN5*, *DPP10*, *HSP90AA1*, HS6ST3, and KCNIP4 for all cell types comparing low vs. high pathology. **(E)** Distributional divergence between low and high pathology groups for the top 2,000 highly variable genes. Each point is one gene, placed by the percentile rank of its Kolmogorov–Smirnov (KS) statistic and Wasserstein distance (WD), computed on log-normalized expression across all cells, including zeros; higher percentiles indicate greater differences between group distributions. Dashed lines mark the 90th percentile. Percentiles are calculated within each analysis’s own set of highly variable genes. Showing analysis from all cell types. Black points are included in the 41 RecurSieve identified gene-panel, with genes *WWOX*, *CNTN5*, *DPP10*, *HSP90AA1*, *HS6ST3*, and *KCNIP4* annotated.

**Figure 4:**
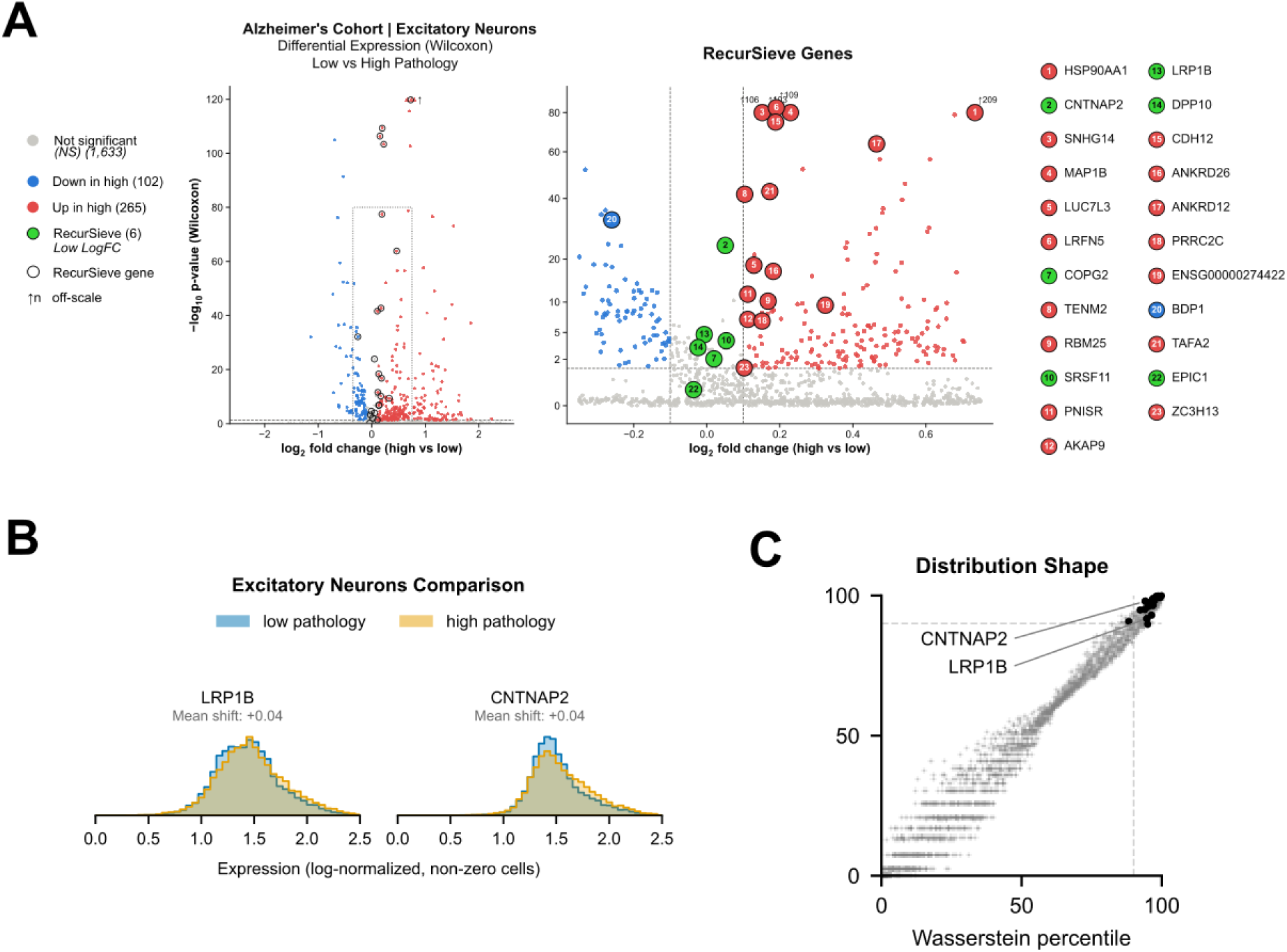
Excitatory Neuron Results in Alzheimer’s Single-cell Data. **(B)** Volcano plot comparing RecurSieve-selected genes (n=23) versus DE-identified genes (Wilcoxon test) in an Alzheimer’s disease single-nucleus cohort in Excitatory neurons only. (Right) ranked list of the 23 RecurSieve-identified genes in the Alzheimer’s cohort, ordered by iteration of selection. **(C)** Excitatory neuron expression distributions (non-zero) and mean shifts of *LRP1B* and *CNTNAP2* comparing low vs. high pathology **(D)** Distributional divergence between low and high pathology groups for the top 2,000 highly variable genes. Each point is one gene, placed by the percentile rank of its Kolmogorov–Smirnov (KS) statistic and Wasserstein distance (WD), computed on log-normalized expression across cells, including zeros; higher percentiles indicate greater differences between group distributions. Dashed lines mark the 90th percentile. Percentiles are calculated within each analysis’s own set of highly variable genes. Showing analysis from excitatory neurons only. Black points are included in the RecurSieve identified gene-panel, with genes *LRP1B* and *CNTNAP2* annotated.

RecurSieve and Wilcoxon DE testing identified both shared and test-specific genes, as visualized on a volcano plot (**Figure 3A and Figure 4A**). In the all cell types analysis, Wilcoxon analysis identified n=103 and n=191 genes that were down- or up-regulated in the high disease state versus low disease state, respectively. RecurSieve analysis identified 41 genes that differed across disease states, including n=19 that were exclusive to RecurSieve and not Wilcoxon analysis, due to subthreshold values for either mean expression change (n=13) or p-value score (n=6)(**Figure 3B**). The remaining 22 genes identified by RecurSieve that met Wilcoxon significance criteria showed moderate to high mean expression fold-changes with statistically significant p-values based on standard testing, with 18 upregulated and 4 downregulated genes in the high pathology group (**Figure 3A,B**). In the Excitatory Neuron analysis,

Wilcoxon analysis identified n=102 and n=265 genes that were down- or up-regulated in the high disease state versus low disease state, respectively. RecurSieve analysis identified 23 genes that differed across disease states, including n=6 that were exclusive to RecurSieve and not Wilcoxon analysis, due to subthreshold values for either mean expression change (n=5) or p-value score (n=1)(**Figure 4A**). The remaining 17 genes identified by RecurSieve that met Wilcoxon significance criteria showed moderate to high mean expression fold-changes with statistically significant p-values based on standard testing, with 16 upregulated and 1 downregulated genes in the high pathology group (**Figure 4A**).

We evaluated how RecurSieve’s iterative recursion contributes to gene selection by comparing to RF important features in the all cell types condition (**Figure 3C**). Notably, RecurSieve identified 33 features not present in RF top features. These, and results for individual cell types, are presented in **Supplemental Table 1**.

We next examined the underpinnings of RecurSieve’s unique testing sensitivity on these empirical data. We first assessed the extent to which genes identified by RecurSieve were driven by expression distribution differences across low and high disease stages (**Figure 3D, 4B**). To do this, we used Kolmogorov-Smirnov (KS) statistic^33^ and Wasserstein distance (WD)^34^ to compare distributional differences for the RecurSieve-identified genes versus all of the 2,000 most variable genes in the analysis (**Figure 3E, 4C**). All highlighted genes ranked above the 92nd percentile demonstrating RecurSieve-identified genes tended to have highly dissimilar expression distributions, consistent with sensitivity revealed by our simulated data (**Figure 2**).

Collectively, these results from empirical data support the basic tenets that RecurSieve can complement standard DE methods by revealing genes with subtle expression differences across groups without overfitting on statistically strong outliers.

To explore whether RecurSieve’s unique testing sensitivity may contribute important biological information, we extended RecurSieve analysis of low versus high disease states across all annotated brain cell populations and compiled a literature-based validation summary for identified genes (**Supplemental Table 1**; **Methods 4.2**). We categorized all genes returned from RecurSieve into four evidence tiers for Alzheimer’s association: High, Medium, Low, and Emerging. This analysis underscored that RecurSieve had recovered expression differences for genes with previous associations to Alzheimer’s disease. We overview example results from all cell types and Excitatory neuron comparisons below, highlighting disease-state expression differences.

As expected by our quantitative summary, a subset of these genes exhibited marked qualitative changes in distribution across low and high disease states. In the all cell types comparison of high and low-pathology groups, we focused on example genes *WWOX, CNTN5, HSP90AA1, HS6ST3, DPP10,* and *KCNIP4* (**Figure 3D**). We focused on *LRP1B* and *CNTNAP2* gene results from Excitatory Neurons (**Figure 4B**).

In results from all cell types, both *WWOX* and *CNTN5* exhibited small log2 fold-change values, yet were prioritized by the algorithm. *WWOX* has been linked to neurodegeneration and tau pathology^35–37^, while gene variants in *CNTN5*, a neuronal adhesion molecule of the contactin family were previously identified as conferring risk for Alzheimer’s through GWAS analysis^38^. Interestingly, *HSP90AA1*, a member of the

Hsp90 chaperone family and implicated in tau stabilization and aggregation^39,40^ was identified as significant by Wilcoxon (log2FC = 0.673; p ≤ 10^-230^), demonstrating RecurSieve’s ability to also recover genes with stronger mean-based differences. Additional genes such as: *KCNIP4* (log2FC=0.016), a calcium-binding Kv channel-interacting protein which is a key resilience factor in early Alzheimer’s^28^; *HS6ST3* (log2FC=−0.034), a member of the heparan sulfate 6-O-sulfotransferase family, whose activity is required for tau uptake^41,42^; and *DPP10* (log2FC=−0.047), a potassium channel partner found in AD tangles^43^, were selected by RecurSieve despite having negligible fold-change in mean-based tests.

Additionally, RecurSieve results from Excitatory neurons (**Figure 4A**) revealed genes such as *LRP1B* (log2FC=−0.008), a low-density lipoprotein (LDL) receptor-related protein that has been shown to lower amyloid-beta production in a non-neuronal cell model expressing an *LRP1B* minireceptor^44^, and *CNTNAP2* (log2FC=0.051), which has been shown to be reduced in the hippocampus of late-onset Alzheimer’s patients^45^. Both genes exhibited low log2FC, highlighting the ability to reveal relevant genes to disease progression.

The ability of RecurSieve to simultaneously recover both subtle and strong signals supported by the literature suggests it is capable of constructing a representative biomarker panel that distinguishes nuanced disease-associated transcriptional changes, ranging from minor regulatory shifts to major pathological drivers.

## 3. Discussion

RecurSieve was developed to expand the sensitivity of detecting genes with complex changes in RNA levels across experimentally-grouped scRNA-seq data. Many standard methods for DE analysis of scRNA-seq data - Wilcoxon rank-sum test, t-test, and logistic regression, for example - were adapted from bulk RNA-seq comparisons and thus rely on univariate mean-based expression comparisons across fixed groups. By iteratively refining cell population labels via GMM clustering, rather than relying on fixed, predefined-labels, RecurSieve allows ‘ground truth’ labels to evolve based on cumulative gene expression and library size. Biologically, this sensitivity arises because molecular cell states are rarely homogeneous across experimental groups and cellular subpopulations may present complex distributional shifts in RNA levels such as bimodality, skewness, and co-expression; group-based RNA expression averaging can mask these effects. In comparing RecurSieve results to those of standard DE methods using staged and empirical scRNA-seq data, we confirmed RecurSieve’s sensitivity and identification of potentially valuable biological information.

Controlled benchmarking using synthetic data highlighted RecurSieve’s sensitivity: while standard methods largely failed to detect staged genes with non-linear changes in RNA levels and little change in average expression, RecurSieve identified ∼20-40% of such genes.

In empirical testing using a large-scale single-nuclei dataset from an Alzheimer’s disease cohort, RecurSieve identified tens of genes with disease stage-specific changes in RNA level distributions with unremarkable changes in average expression that were missed by standard Wilcoxon testing. A systematic survey of the literature revealed that many of these genes - including *WWOX*, *CNTN5*, *DPP10*, *LRP1B*, and *CNTNAP2*, had been previously associated with Alzheimer’s disease processes. RecurSieve also identified large-effect expression changes in genes like *HSP90AA1* that were also identified in standard testing, demonstrating some shared sensitivity with standard DE methods and discoverability spanning a spectrum of significant transcriptional changes. In the context of human disease, application of RecurSieve alongside standard methods could serve to expand panels of biologically-relevant genes, including as potential biomarkers.

RecurSieve also offers practical advantages in computational interpretability. The algorithm converged rapidly on large datasets (typically 15-50 iterations). Unlike less-interpretable deep learning approaches^46^, RecurSieve provides a transparent, iterative feature selection process where each gene’s contribution to label refinement can be tracked, offering a clear rationale and interpretable history for genes included in the final signature.

Several limitations warrant consideration. First, RecurSieve relies on the assumption that the condition-varying transcriptional signals can be approximated by a mixture of Gaussians; highly complex, multi-modal distributions with overlapping components may challenge the GMM clustering and potentially erode the reliability of iterative label assignment. Second, while the nature of the algorithm allows for the potential discovery of non-linear patterns, it is computationally more intensive than a single-pass differential expression test. In practice, implementing RecurSieve remains tractable for modern single-cell RNA-seq datasets (**Figure 1B**). Third, the current implementation focuses on binary classification across two groups; extending the RecurSieve framework to multi-class comparisons (e.g., multiple Braak stages concurrently, rather than high and low pathology) would require adapting the GMM component to handle multiple clusters and might necessitate more complex iteration-stopping criteria. Fourth, while we benchmarked RecurSieve against standard DE methods, we did not compare it against methods designed to detect differential distributions or single-cell-specific feature selection frameworks such as variational autoencoder-based approaches (e.g., scVI^20,47^, scANVI^47^), Donor aware methods, or deep learning wrappers (e.g., scGen^48^). Consequently, while RecurSieve offers extended and complementary sensitivity as compared to traditional univariate tests, its relative performance against these model-based selectors remains to be determined and distribution focused tests in particular are likely to outperform RecurSieve for single-gene shape changes. Fifth, training and testing splits were performed at the cell level. This was a deliberate choice, as a donor-level split would leave few donors per group in the test set. Because cells from the same donor may appear in both sets, classification metrics may be inflated and should be interpreted as a measure to distinguish labels rather than generalization to new donors. Lastly, RecurSieve includes a null permutation test as a robustness check that determines statistical significance and seed stability for individual genes, a process that can be computationally resource-intensive. Additionally, we recommend that the biological relevance of each selected gene be confirmed through experimental validation, manual literature review and by running standard DE tests specifically on the identified panel to supplement statistical metrics.

Future improvements could focus on optimizing computational efficiency of the recursive loop, potentially through parallelization of RF training or approximation techniques for GMM fitting. Additional avenues of exploration include integration of Large Language Models (LLMs) to automate initial literature validation of low-fold-change candidates, which could streamline manual curation for confirming biological relevance. Furthermore, benchmarking alternative feature selection methods such as mutual information, SVMs, or gradient boosting in place of the current classifier could determine whether different importance metrics yield more robust or biologically specific gene panels.

In conclusion, RecurSieve offers a unique approach towards single-cell feature selection by moving beyond the limitations of mean-based DE testing approaches. By embracing the complexity of gene expression distributions, RecurSieve reveals subtle yet robust alterations in transcriptional signals that may help achieve a more complete understanding of the molecular variability present in complex tissue in health and disease.

## 4. Methods

### 4.1.1 RecurSieve Software Architecture

The package is constructed as an object-oriented architecture. The main class ‘recursievè can be instantiated minimally with parameters defining an AnnData^49^ or AnnSQL^50^ object, the two comparison groups found in the ‘obs’ layer of the AnnData object (‘group1’, ‘group2’), and ‘field_namè to define the ‘obs’ label field of the groups. After light preprocessing (filter, normalize, transform), the dataset is split into training (70%) and testing (30%) for training and assessment.

### 4.1.2 Algorithm Description

RecurSieve distinguishes between two groups, specifically in single-cell datasets. The algorithm initially approaches the two defined groups as a classification problem using RandomForestClassifier via Scikit-learn^51^, then feature selection based on the trained model is calculated using Gini Impurity^29^. Next, the top feature (gene) by importance score is saved for use along with the total UMI counts of each single-cell library to cluster two new group labels using a Gaussian Mixture Model (GMM). These labels are then used to train a subsequent RandomForestClassifier with all features (genes) minus the initial top-ranking important feature. Next, Gini Impurity is calculated on the new model, and the highest-ranking gene is saved. The mean of both genes is calculated, then used along with the total UMI counts of each single-cell library to establish two new training labels using a GMM. This process of generating dynamic training targets and identifying important gene features via RF modeling will continue until GMM training labels stabilize, with each gene being saved at each iteration. Formally, we define the algorithm as the following iterative procedure. Let 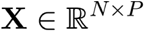 represent the normalized gene expression matrix for *N* cells and *P* genes, and let 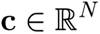 denote the vector of total UMI counts (log1p-transformed). We initialize the label vector *y^(0)^*with the original group assignments and the set of available genes 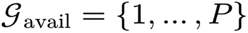 . At each iteration 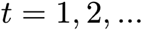 :

1. **Gene Selection:** A RF classifier *F_t_* is trained on the current data 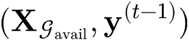 . The importance score *I_j_* for each gene 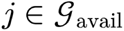 is computed based on the total reduction in Gini impurity across all decision trees. The top gene *g_t_* is selected as:

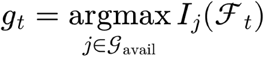

The selected gene is removed from the available pool: 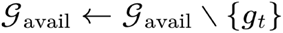 .
2. **Label Update**: To generate the training targets for the next iteration, we construct a feature matrix *Z^(t)^* combining the total UMI counts *c* and the aggregated expression of all selected genes 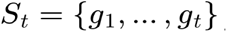 . The aggregation 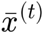 is defined as the mean expression across the selected panel:

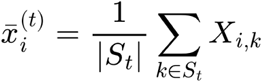

A two-component Gaussian Mixture Model (GMM) is fitted to the rows of 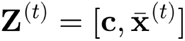. The new label vector *y^(t)^* is assigned such that 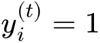 if cell *i* belongs to the GMM component with the higher mean expression, and 0 if otherwise.
**3. Termination:** The algorithm terminates when the label vector stabilizes **(Methods 4.1.3)**.

### 4.1.3 Assessment of Recursive iterations and Gene Panel

After each recursive iteration, RecurSieve re-clusters the cells using the current set of selected genes and compares the binary GMM labels to the previous iteration. To account for possible label swapping between the two classes, we compute the minimum label disagreement rate between the current and previous assignments. Let *D_t_* denote the minimum label disagreement rate between the binary GMM label assignments at iterations *t* and *t*-1, accounting for possible swapping of the two class labels. We define:

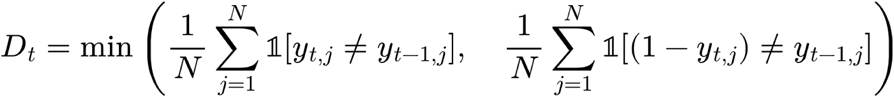

where *y_tj_* is the binary cluster label for cell *j* at iteration *t*. The corresponding convergence stability is then

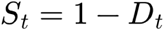

which measures the fraction of labels that remain unchanged. Recursion continues until *D_t_* falls below a predefined threshold (Default: 0.01), indicating that the labels have stabilized, or until no improvement is made after 10 iterations (Patience: 10). Additionally, repeating two cycles of oscillation with exact label assignments triggers early termination. To examine the quality of the model and the gene panel, we include two metrics: (1) the seed stability metric can be calculated using the ‘seed_stability’ method, which repeats the algorithm (Default: 10) while changing the random seed metric. Examining the returned ‘gene_frequency’ result can be used to measure the gene selection stability with respect to the seed. (2) The ‘coexpression_null’ method is a permutation test that refits RecurSieve on randomized datasets (default: 100). In each one, genes and the group labels are permuted. Each gene’s p-value is the fraction of randomized panels that also select it, estimated as (1 + n)/(1 + N), where n is the number of randomized panels that include the gene and N is the number of shuffles. P-values are then adjusted for multiple testing using the Benjamini-Hochberg (BH) procedure for all genes in the final panel. Lastly, we maintain a log of AUC, OOB, and accuracy to evaluate algorithm performance at each stage of its recursion. These metrics are calculated using the Sklearn metrics class methods: accuracy_score and roc_auc_score.

### 4.1.4 Model Parameters

Each RF model maintains stable parameters through all iterations. The default parameters are as follows: n_estimators: 300, max_depth: 10, min_samples_split: 2, min_samples_leaf: 1, class_weight: balanced_subsample. The GMM parameters in each iteration are all default, with n_components set at 2. Verbose model parameters for plotting, summary, seeds, and more are provided at the package repository https://github.com/ArpiarSaundersLab/RecurSieve

### 4.2 Public Alzheimer’s Dataset

We utilized a publicly available, published single-nuclei gene expression (snRNA-seq) dataset from Alzheimer’s patients^28^. The dataset consisted of 424,528 nuclei derived from three key cortical regions of the brain: the Prefrontal, Precuneus, and Primary Visual cortices. The data includes 46 donors stratified by disease severity, with 18 exhibiting no or low AD pathology (Braak stages 0-II), 10 with intermediate pathology (Braak stages III-IV), and 18 with high AD pathology (Braak stages V-VI). This dataset enables granular analysis across clinical and biological features, including Braak stages, age, amyloid levels, brain region, ethnicity, and sex. Samples were sequenced using Drop-seq or 10x Genomics platforms, with primary visual cortex samples predominantly generated by Drop-seq. For our analysis, we focused on all cell types combined and Excitatory neurons, while utilizing the high- and low-disease pathology metadata included.

#### 4.2.1 Default RecurSieve Alzheimer’s Dataset Preprocessing Pipeline

The RecurSieve package expects a raw AnnData or AnnSQL layer. The raw layer of the dataset is then filtered to contain only high and low disease pathology cells. Once filtered, the dataset was downsized to 50,000 cells for training and testing using the Scanpy ‘samplè method. During preprocessing, all mitochondrial, ribosomal, and hemoglobin genes, and cells with few or rare genes were removed. The dataset was then normalized, log1p-scaled, and subset to the top 2000 most variable genes using the appropriate Scanpy methods from within the RecurSieve class.

#### 4.2.2 Alzheimer’s Dataset Analysis

Scanpy differential expression was performed for the two groups compared by RecurSieve (high/low pathology) using the Wilcoxon rank-sum test^9^, which is a standard differential expression method included in the Scanpy suite of analysis tools. We then compared the list of genes identified by DE to the list identified by RecurSieve using custom scripting. Lastly, all DE and RecurSieve-found genes are presented in **Supplemental Table 1** (by cell type) and were binned by literature support for Alzheimer’s disease: High (genetic or causal evidence), Medium (direct pathology evidence), Emerging (single or limited studies), and Low (no disease-specific literature). Seed stability (10 seeds) and Benjamini–Hochberg-adjusted permutation p-values (**100 permutations; Methods 4.1.3**) are reported for each RecurSieve gene in **Supplemental Table 1**.

### 4.3 Simulations

We constructed simulated datasets using the Python implementation of Splatter^30^. Within these datasets, we introduced staged gene expression relationships across 10 specific genes by substituting simulated counts with bimodal, trimodal, skew, and coexpression patterns for one condition. For the bimodal scenario, each target gene in group 1 followed a Normal distribution centered at *µ*. Cells in group 2 were split into two equal subpopulations: one drawn from a low component (*µ*-*δ*) and the other from a high component (*µ*+*δ*). The separation parameter δ was determined to ensure the mixture mean and median of group 2 matched group 1 exactly, isolating differences to distributional shape rather than central tendency. Next, we created a trimodal condition where group 2 cells were distributed across three modes (*µ*-*δ,µ,µ*+*δ*) with equal probability, again preserving the global mean and median of group 1. We further implemented a skew scenario where group 1 followed a right-skewed skew-normal distribution and group 2 followed a mirrored left-skewed counterpart; the location parameters were shifted to align the means despite opposing tail directions. Finally, for the coexpression benchmark, we maintained identical marginal distributions for both groups but induced positive pairwise correlation among the staged genes exclusively within group 2, testing dependence-only recovery. All other genes in the simulated datasets retained standard simulation parameters, creating a background of conventional differential and non-differential signals to benchmark recovery of the staged set, and were composed of 2000 cells by 2000 genes. Wilcoxon and t-test genes were retained at Benjamini–Hochberg adjusted p < 0.05, and logistic regression, which does not produce p-values, was represented by its top 50 genes by score. For each method, recovery was measured as the number of the 10 staged genes present in its selected gene set, compared against the final RecurSieve panel.

### 4.4 Software Packages Integrated in RecurSieve

Python (3.12) packages implemented as part of RecurSieve include: numpy (v2.3.5), pandas (v2.3.3), scanpy (v1.11.5), anndata (v0.12.7), annsql (v1.0.3), seaborn (v0.13.2), matplotlib (v3.10.8), scikit-learn (v1.8.0), and scipy (v1.16.3).

## 5. Declaration of Generative AI and AI-Assisted Technologies

During the preparation of this work, the authors used Claude Opus 5 for debugging code errors, drafting analysis code, documentation, and descriptive text. Proton Lumo(v2) was also used for editing and proofreading of manuscript text. After using these tools, the authors reviewed and edited the content and take full responsibility.

## 6. Operation

RecurSieve requires Python [3.12 or later] and runs on macOS, Linux, and Windows. It was tested on a MacBook with an 8-core CPU (16 GB RAM) (**Figure 1B**) and a Ubuntu 24.04 laptop with a 12-core CPU (i7-1255U; 40GB RAM). Dependencies (**Methods 4.4**) are installed automatically:

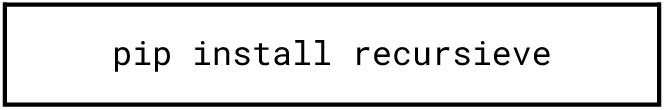

A minimal run takes an AnnData object, the metadata field containing group labels, and the two groups to compare:

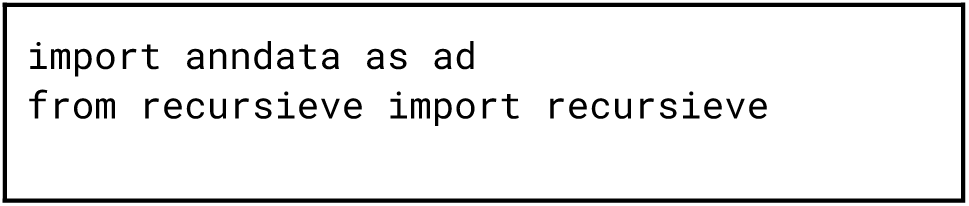

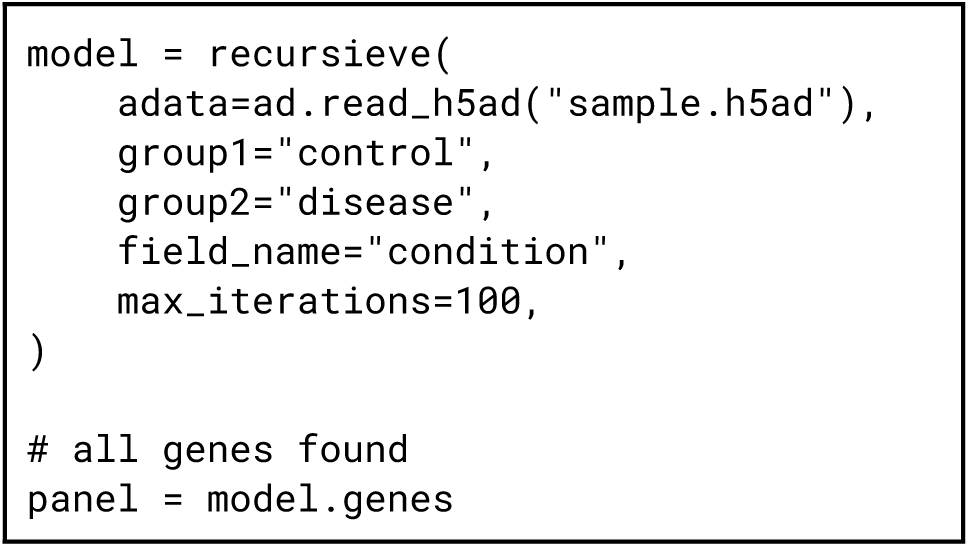

Full parameters and examples are available at https://github.com/ArpiarSaundersLab/RecurSieve.

## 7. Software Availability

Software available from: https://github.com/ArpiarSaundersLab/RecurSieve

Package available from: https://pypi.org/project/recursieve/

Source code available from: https://github.com/ArpiarSaundersLab/RecurSieve

Archived source code: https://doi.org/10.5281/zenodo.24016571

License: MIT

## 8. Data Availability

### Underlying data

NCBI Gene Expression Omnibus: Molecular Signatures of Resilience to Alzheimer’s Disease in Neocortical Layer 4 Neurons, Accession GSE263468. https://www.ncbi.nlm.nih.gov/geo/query/acc.cgi?acc=GSE263468

Simulated datasets can be regenerated with the scripts included in the archived source code https://doi.org/10.5281/zenodo.24016571

### Extended data

Zenodo: Extended data for: RecurSieve: A Python Package Using Iterative Machine Learning to Detect Subtle Gene Expression Patterns in Single-Cell Data.

https://doi.org/10.5281/zenodo.24015171

This project contains the following extended data:

- **Supplemental_Table_1.xlsx**: RecurSieve genes for all cell types combined and each cell type individually, with Alzheimer’s disease literature evidence tiers.

Data available under terms of the Creative Commons Attribution 4.0 International license (CC-BY 4.0).

## 10. Author Contributions

Conceptualization: A.M., K.P., L.BA.

Methodology: A.M., K.P.

Software: A.M., K.P.

Validation: A.M., K.P.

Formal analysis: A.M., K.P.

Investigation: A.M., K.P., A.S., G.C, C.F.

Data curation: A.M., K.P.

Writing original draft: A.M., K.P.

Writing review & editing: A.M., K.P., A.S., L.BA, G.C, C.F.

Visualization: A.M., K.P.

Supervision: K.P., A.S.

Project administration: K.P., A.S.

Funding acquisition: A.S.

